# Identification of Lysine Isobutyrylation as A New Histone Modification Mark

**DOI:** 10.1101/2020.08.31.270165

**Authors:** Zhesi Zhu, Zhen Han, Levon Halabelian, Xiangkun Yang, Jun Ding, Nawei Zhang, Liza Ngo, Jiabao Song, Hong Zeng, Maomao He, Yingming Zhao, Cheryl H. Arrowsmith, Minkui Luo, Michael G. Bartlett, Y. George Zheng

**Affiliations:** Department of Pharmaceutical and Biomedical Sciences, College of Pharmacy, University of Georgia, Athens, GA 30602, United States; Structural Genomics Consortium, University of Toronto, Toronto, ON, M5G 1L7, Canada; Department of Medical Biophysics, University of Toronto, Toronto, ON, M5G 1L7, Canada; Princess Margaret Cancer Centre, University Health Network, Toronto, ON, M5G 2M9, Canada; Ben May Department for Cancer Research, The University of Chicago, Chicago, IL 60637, United States; Chemical Biology Program, Memorial Sloan Kettering Cancer Center, New York, NY 10065, United States; Program of Pharmacology, Weill Cornell Medical College of Cornell University, New York, NY 20021, United States

**Keywords:** lysine isobutyrylation, isobutyrate, valine, HAT, KAT, epigenetics, histone modification

## Abstract

Short-chain acylation of lysine residues in eukaryotic proteins are recognized as essential posttranslational chemical modifications (PTMs) that regulate cellular processes from transcription, cell cycle, metabolism, to signal transduction. Lysine butyrylation was initially discovered as a normal straight chain butyrylation (Knbu). Here we report its structural isomer, branched chain butyrylation, i.e. lysine isobutyrylation (Kibu), existing as a new PTM on nuclear histones. Uniquely, isobutyryl-CoA is derived from valine catabolism and branched chain fatty acid oxidation which is distinct from the metabolism of n-butyryl-CoA. Several histone acetyltransferases were found to possess lysine isobutyryltransferase activity, especially p300 and HAT1. We resolved the X-ray crystal structures of HAT1 in complex with isobutyryl-CoA that gleaned an atomic level insight into HAT-catalyzed isobutyrylation. RNA-Seq profiling revealed that isobutyrate greatly affected the expression of genes associated with many pivotal biological pathways. Our findings identify Kibu as a novel chemical modification mark in histones and suggest its extensive role in regulating epigenetics and cellular physiology.

## Introduction

Protein posttranslational modifications (PTMs) fundamentally impact on cellular physiology and phenotype in eukaryotic organisms.^1^ Fatty acylation on the side-chain amino group of lysine residues has been recognized as an important type of reversible PTM marks that impart various regulatory functions on key cellular processes such as gene transcription, metabolism, protein homeostasis, and signal transduction.^2–4^. So far about 20 types of lysine acylations have been discovered including acetylation, propionylation, butyrylation, crotonylation, succinylation, malonylation, and glutarylation.^5–8^ Lysine acetyltransferases (KATs) are the writer enzymes that introduce acylation marks on specific lysine residues using acyl-CoA molecules as the acyl donor.^9^ Acyllysines are recognized by downstream reader proteins, and enzymatically reversed by eraser proteins, histone deacylases (HDAC).^10,11^ Deregulation of lysine acylation dynamics caused by aberrant expression or mutation of either writer, reader, or eraser proteins is broadly associated with various disease phenotypes including inflammation, neurodegeneration, cancer, etc.^12–14^ Even occurring on the same residues, different acylations could result in distinct biological outcomes. For instance, butyrylation competes with acetylation on H4K5/K8 and prevents the binding of the reader protein Brdt on these loci, which causes delayed histone removal and gene expression in spermatogenic cells.^15^ It remains an imperative task to map out cellular substrates and modification sites of different lysine acylations and investigate their functional impacts in different physiological and pathological pathways.

In the last decade, the development of high resolution mass spectrometry has greatly facilitated discovery of many novel lysine acylation marks.^16,17^ Lysine butyrylation (Kbu) was first discovered by Zhao and coworkers as a normal straight chain n-butyrylation (Knbu) which is biochemically dependent on n-butyryl-CoA.^18^ Recent studies demonstrated that KAT members p300/CBP possess lysine n-butyryltransferase activity.^19^ Etiologically, n-butyryl-CoA is a metabolic intermediate from the fatty acid oxidation pathway and therefore, serving as an ample donor source for Knbu. From a chemical perspective, butyrylation mark may also exist in its isomeric structure, i.e., isobutyrylation (2-methylpropionylation) **(Figure 1)**. Our proposal is further inspired by the natural existence of isobutyryl-CoA which is endogenously generated as a result of valine catabolism and the oxidation of branched chain fatty acids.^20^ Taking into account the physiological existence of isobutyryl-CoA and its structural relevance with n-butyryl-CoA, we posit that isobutyryl-CoA may also function as an acyl group donor and lead to an unexplored acylation mark in proteins, lysine isobutyrylation. Pathologically, dysregulation of isobutyryl-CoA level caused by genetic deficiency of isobutyryl-CoA dehydrogenase (IBD) is associated with multiple disease symptoms including speech delay, anemia, and dilated cardiomyopathy in newborn patients while the mechanism underlying these disorders are still poorly defined.^21,22^ Hence, understanding the catabolism of isobutyryl-CoA in relationship with dynamic regulation of protein function through the PTM mechanism has a profound pathophysiological significance. In the present study, we report lysine isobutyrylation (Kibu) as a *bona fide* PTM in nuclear histones throught a combined suite of biochemical, biophysical, and cellular studies.

**Figure 1.**
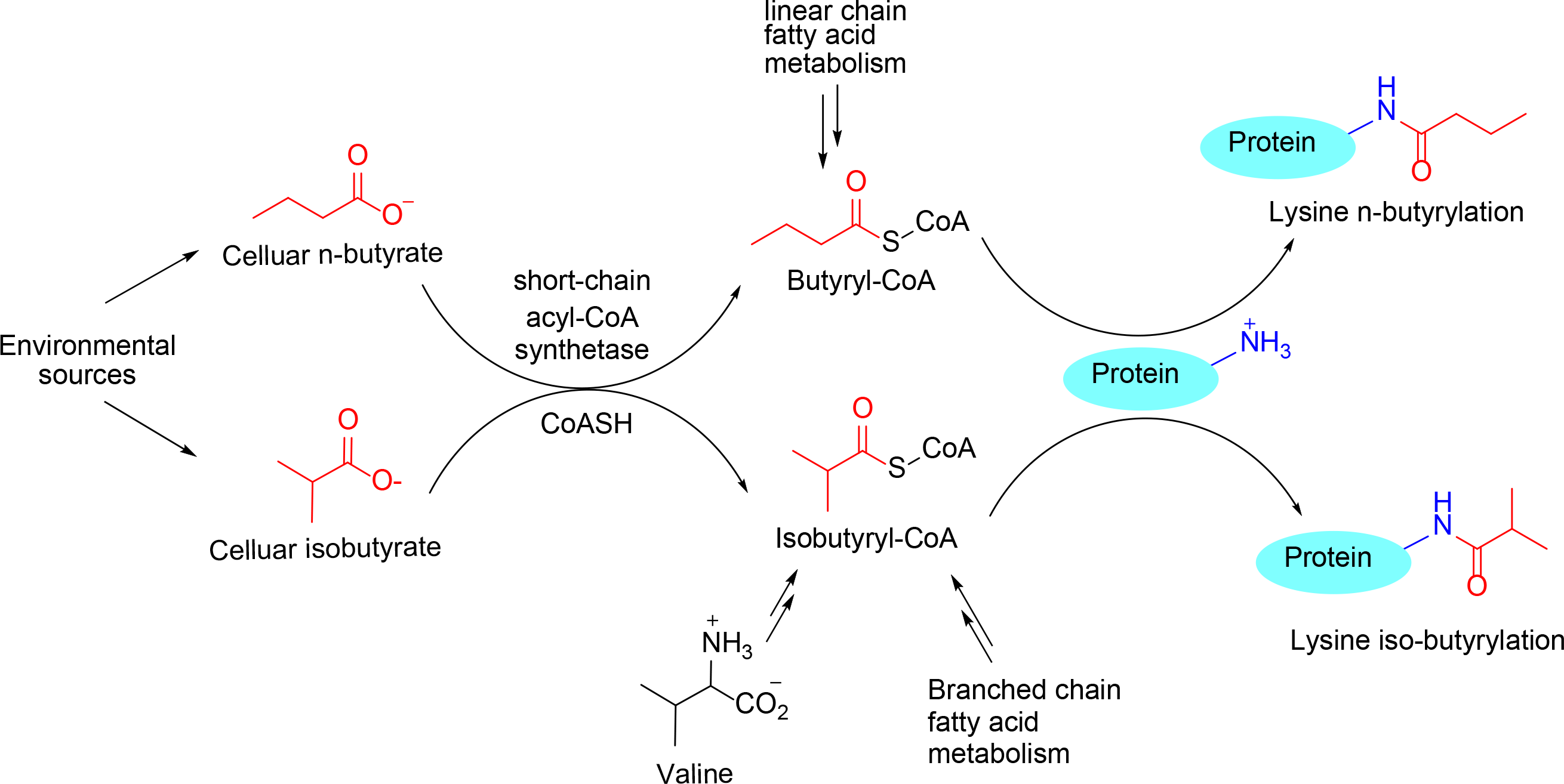
Distinct pathways of two isoforms of butyryl-CoA and isobutyl-CoA and their consequence in leading to protein lysine butyrylation. Lysine butyrylation was identified in 2007^18^ and was recognized as n-butyrylation. In this study, we found that lysine butyrylation also exists in the form of isobutyrylation in histones. Endogenous n-butyryl-CoA is derived from fatty acid metabolism while isobutyryl-CoA is derived from valine metabolic pathway. Exogenous butyrate and isobutyrate can be converted to their acyl-CoA by the action of cellular short-chain acyl-CoA synthetases.

## Results

### Isobutyryl-CoA is an abundant metabolite in the mammalian cell

Upon its discovery with mass spectrometry, lysine butyrylation (Kbu) was defined as a straight chain n-butyrylation.^18^ Nevertheless, two isomeric structures may contribute to the same molecular mass of the butyryl mark (+70 au): either normal linear butyryl or branched isobutyryl group. Lysine acylations rely on acyl-CoA molecules as the reactive acyl group donor **(Figure 1)**; therefore, an ample pool of acyl-CoA is a prerequisite for the incidence of cognate lysine acylation. To determine cellular butyryl-CoA composition, we measured the levels of n- and iso-butyryl-CoA in human embryonic kidney 293T (HEK293T) cells with high-performance liquid chromatography tandem mass spectrometry (HPLC-MS/MS). Iso- and n-butyryl-CoA standards were resolved on the chromatogram at retention times of ~13.85 and 14.05 minutes respectively, and monitored at the same ion transition (838→331) on the mass spectrometer **(Figure 2A)**. The chromatographic peak area of n- or iso-butyryl-CoA molecules was integrated as the quantitation reference. Then HEK293T cell’s acyl-CoA extracts were subjected to the same HPLC-MS/MS analysis and the abundance of n- and iso-butyryl-CoAs was compared. Duplicate experiments were conducted, showing that the ratios of iso- and n-butyryl-CoA were 2~3:1 **(Figure 2B)**. This data revealed that the cellular butyryl-CoA is a mixture of two isomers, both of which may act as acyl donors leading to lysine butyrylation on cellular proteins.

**Figure 2.**
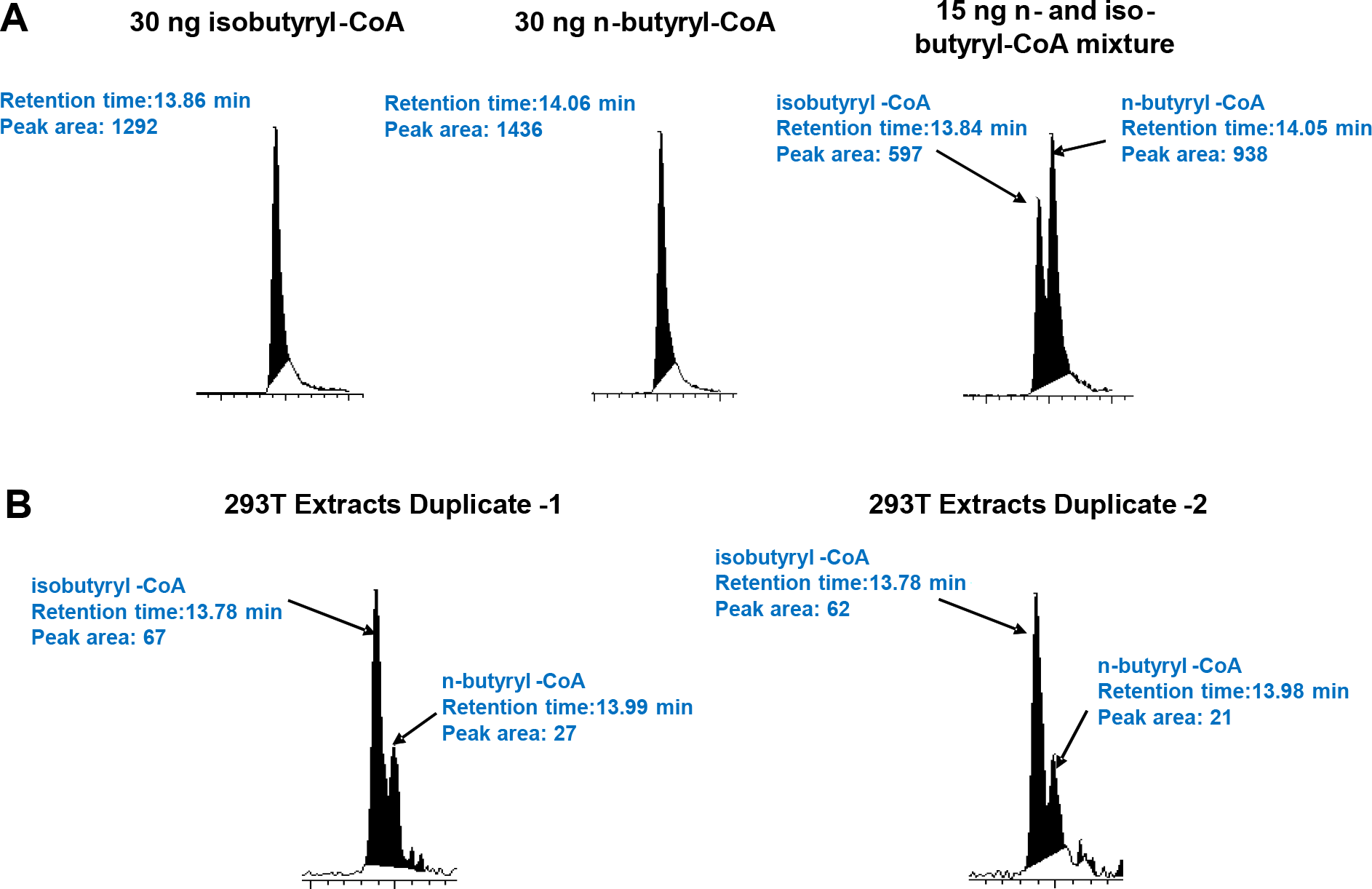
Detection and quantification of n- and iso-butyryl-CoA using LC-MS/MS. **A.** Control experiments showing that N- and iso-butyryl-CoA were separated with the LC-MS/MS system with the retention times at 14.06 and 13.86 minutes and were detected at the same ion transition 810 → 313. **B.** n- and iso-butyryl-CoA from the 293T cell extracts were detected under the same LC-MS/MS condition. Their levels were compared by using the integrated chromatographic peak areas.

### Cellular metabolism of isobutyrate and valine lead to isobutyryl-CoA production

Acyl-CoA molecules are metabolic intermediates mostly from glucose catabolism, fatty acid β-oxidation, and amino acid degradation.^23^ Also short chain fatty acids such as propionate and butyrate taken up from surrounding media can be converted to cognate acyl-CoA molecules with the catalysis of short chain acyl-CoA synthetases (ACS).^24,25^ It is thus highly possible that isobutyryl-CoA can be synthesized from this pathway with isobutyrate as the source agent. Importantly, isobutyryl-CoA is a key intermediate of valine metabolism pathway so valine may also stimulate biosynthesis of isobutyryl-CoA **(Figure 1).**^26^ These isobutyryl-CoA synthetic pathways would further strengthen the possibility that isobutyryl-CoA acts as an important regulator in physiological processes. Herein, we studied the effect of isobutyrate and valine treatment on the change of cellular isobutyryl-CoA level. 293T cells were treated with varied concentrations of deuterated sodium isobutyrate (d7-isobutyrate), and acyl-CoA molecules were extracted and analyzed with the HPLC-MS/MS method aforementioned. The levels of butyryl-CoA molecules were measured and normalized to total cellular proteins **(Supplementary Figure S1A)**. With the increase of d7-isobutyrate doses, d7-isobutyryl-CoA level increased drastically until a plateau was reached at 5 mM of d7-isobutyrate. The level of endogenous, non-isotopic isobutyryl-CoA remained barely changed. We also tested how isobutyryl-CoA level would change in response to valine feeding. The data showed that treatment of 293T cells with valine significantly increased the level of isobutyryl-CoA until a plateau was reached at the concentration of 5 mM valine **(Supplementary Figure S1B)**. In contrast, the level of n-butyryl-CoA was not affected by valine treatment. This observation is consistent with the aforementioned valine metabolic pathway, in which isobutyryl-CoA is an important intermediate.^26^ Taken together, these data demonstrate that isobutyryl-CoA can be biosynthesized from the acyl-CoA synthetase pathway and valine metabolism, which not only defines the anabolism of isobutyryl-CoA in mammalian cells but also provides us with an amenable approach to study the dynamic change of lysine isobutyrylation levels in cellular proteins.

### KAT enzymes catalyze lysine isobutyrylation

Lysine acylations are enzymatically driven by KATs, which can covalently deposit acyl groups from the cosubstrate acyl-CoA to lysine residues in protein substrates. Identification of isobutyryl-CoA as an ample acyl-CoA donor points out the high probability for the existence of lysine isobutyrylation in cellular proteins. In this regard, it is necessary to examine whether and which KATs can catalyze this acylation reaction. We quantitatively measured the acylation activities of nine human KATs from three major KAT families, MYST, GCN5/PCAF/HAT1, and p300/CBP, using a fluorogenic assay that was designed to quantify the byproduct CoA.^27^ Histone peptides H3(1-20) or H4(1-20) was used as the acyl acceptor to characterize the acetyl-, propionyl-, n-butyryl-, and isobutyrylation activities of individual KAT enzymes **(Figure 3)**. Consistent with our recent study,^28^ all the tested KAT enzymes show remarkable Kpr activity, more than 20% of their nascent acetyl transfer activity, which demonstrates that almost all eukaryotic KATs may possess this intrinsic activity. Nevertheless, further increase of acyl chain length to four carbon units leads to a drastic decrease of butyrylation activity (Kbu) of KAT enzymes. Among the tested KAT enzymes, HAT1 showed outstanding isobutyrylation (Kibu) activity with ~25% of its acetylation activity, followed by HBO1 while the other KAT enzymes exhibited much lower or even barely detectable Kibu activity. p300 is known for its great cofactor promiscuity, functioning as lysine acetyl-, propionyl-, butyryl-, and crotonyltransferase.^2,18,19^ Its unique acyl-CoA binding pocket enables p300 to bind with larger acyl-CoA molecules without need of protein engineering.^29,30^ Therefore, we carried out steady-state kinetic characterization of p300 and HAT1 with various acyl-CoAs to determine their acyltransferase activities. The results show that the catalytic specificity constants *k_cat_/K_m_* of p300 with n- and isobutyrylation activities were 0.25 min^-1^.μ^-1^ and 0.13 min^-1^.μ^-1^ respectively, about 7% and 13% of p300 acetylation activity **(Figure 4)**. In comparison, Knbu and Kibu activity of HAT1 was about 13% and 31% of its acetylation activity, respectively. Thus, both HAT1 and p300 possessed appreciable activities of lysine n-butyrylation and iso-butyrylation.

**Figure 3.**
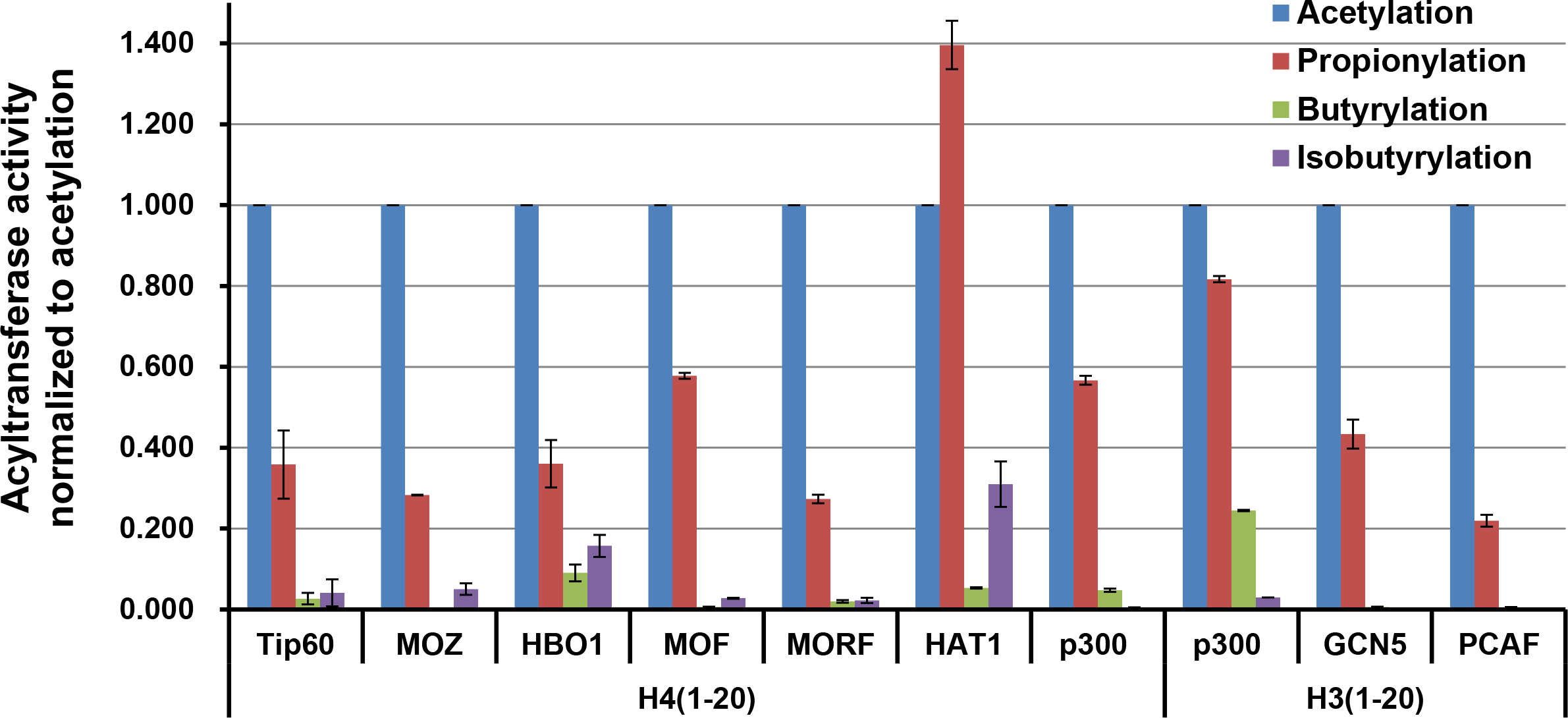
Measurement of lysine acylation activity of KAT enzymes. Lysine acetyl-, propionyl-, n-butyryl-, and isobutyryltransferase activity of each KAT enzymes was tested using a fluorometric CPM assay, with fixed concentration of acyl-CoAs and H3(1-20) and H4(1-20) histone peptide substrates. All tested KATs show strong activity on lysine acetylation and appreciable activity on lysine propionylation. HAT1 shows the strongest activity of carrying out lysine isobutyrylation, about 25% of its acetyltransferase activity.

**Figure 4.**
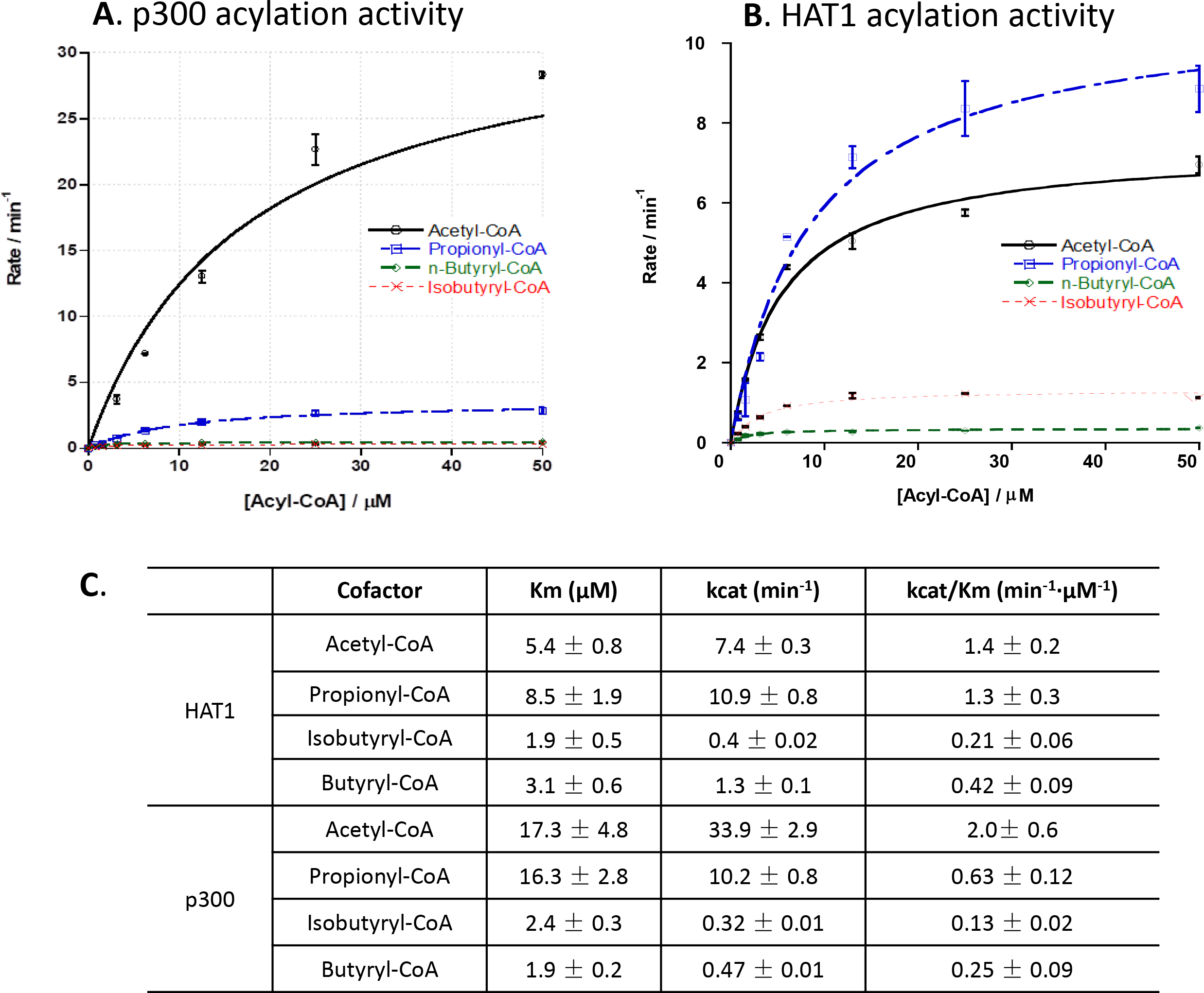
Kinetic characterization of p300 and HAT1 acylation activities. p300 or HAT1 was incubated with individual acyl-CoA molecule at varied concentrations and H4(1-20) peptide substrates. The enzymatic reaction was quantified using CPM assay. The reaction rate-acyl-CoA concentration were plotted with the Michaelis-Menten equation to get the kinetic constants *K_m_* and *k_cat_*. The *k_cat_/K_m_* value were used to evaluate lysine acylation activity. **A.** initial velocity curves of p300 with acetyl-, propionyl-, n-butyryl-, and isobutyryl-CoA. **B.** initial velocity curves of HAT1 with acetyl-, propionyl-, n-butyryl-, and isobutyryl-CoA. **C.** Kinetic parameters from the velocity data fitting. Experimental conditions: 200 μM H4(1-20) peptide, 0-50 μM acyl-CoA, 40 nM or 100 nM HAT1, 20 nM or 100 nM p300, 15 min reaction time, temperature at 30 °C.

HAT1 and p300 isobutyryltransferase activity on synthetic H3 and H4 peptides was further confirmed by MALDI-MS analysis of the reaction mixtures (**Supplementary Figure S2**). The recombinant p300 and HAT1 were incubated with isobutyryl-CoA and H3(1-20) (for p300) or H4(1-20) peptides (for HAT1) for 1 h. Next, the reaction mixture was subjected to MALDI-MS test. Product peaks (M+70) were observed in both p300 and HAT1 catalytic reactions, demonstrating that both enzymes can catalyze isobutyrylation on peptide substrates. We further tested if the isobutyrylation activity of HAT1 on histone H4 substrate can be detected with Western blot using the commercially available anti-butyryllysine antibody (PTM Biolab, Cat# PTM-301). Although this antibody was designed for detection of Knbu,^15,31^ incubation of n- or iso-butyryl-CoA, histone H4 and HAT1 drastically increased the band intensity of both n-butyrylated H4 and iso-butyrylated H4, whereas lack of HAT1 induced little histone labeling (**Supplementary Figure S3**). Hence, this anti-Knbu antibody was also able to recognize Kibu mark. Our finding on the promiscuous specificity of the anti-butyryllysine antibody indicates that some butyrylated lysines identified in previous work could be a mixture of n- and iso-butyrylated lysines. Overall, the biochemical measurements and western blot data validated the Kibu activity of HAT1 and p300. Importantly, the capability of PTM-301 antibody in recognizing Kibu mark allows for a technical means to study protein Kibu signals from the cellular contexts.

### Lysine isobutyrylation is a bona fide PTM mark on nucleosomal histones

We next focused on the detection of lysine isobutyrylation (Kibu) on cellular proteins. 293T cells were treated with sodium d7-isobutyrate for 16 h, followed by extraction of the nuclear histone proteins or whole cellular proteome. The extracted proteins were resolved on a SDS-PAGE gel and analyzed with Western blot using the anti-Kbu antibody PTM-301. The chemiluminescent protein bands on the Western blot membrane are collective signals corresponding to Knbu and Kibu levels because of the promiscuity of this antibody. Nevertheless, any intensity change upon d7-isobutyrate treatment will reflect changes in the Kibu level because isobutyrate treatment induced synthesis of isobutyryl-CoA rather than n-butyryl-CoA. As shown in **Figure 5**, both whole proteome extracts and histone extracts showed increased Kibu levels on histones H3 and H4, as a result of isobutyrate treatment. Thus, Kibu is a *bona fide* histone PTM and is driven by isobutyryl-CoA. Surprisingly, under our condition, no appreciable change of chemiluminescence intensity was observed on non-histone proteins upon isobutyrate treatment.

**Figure 5.**
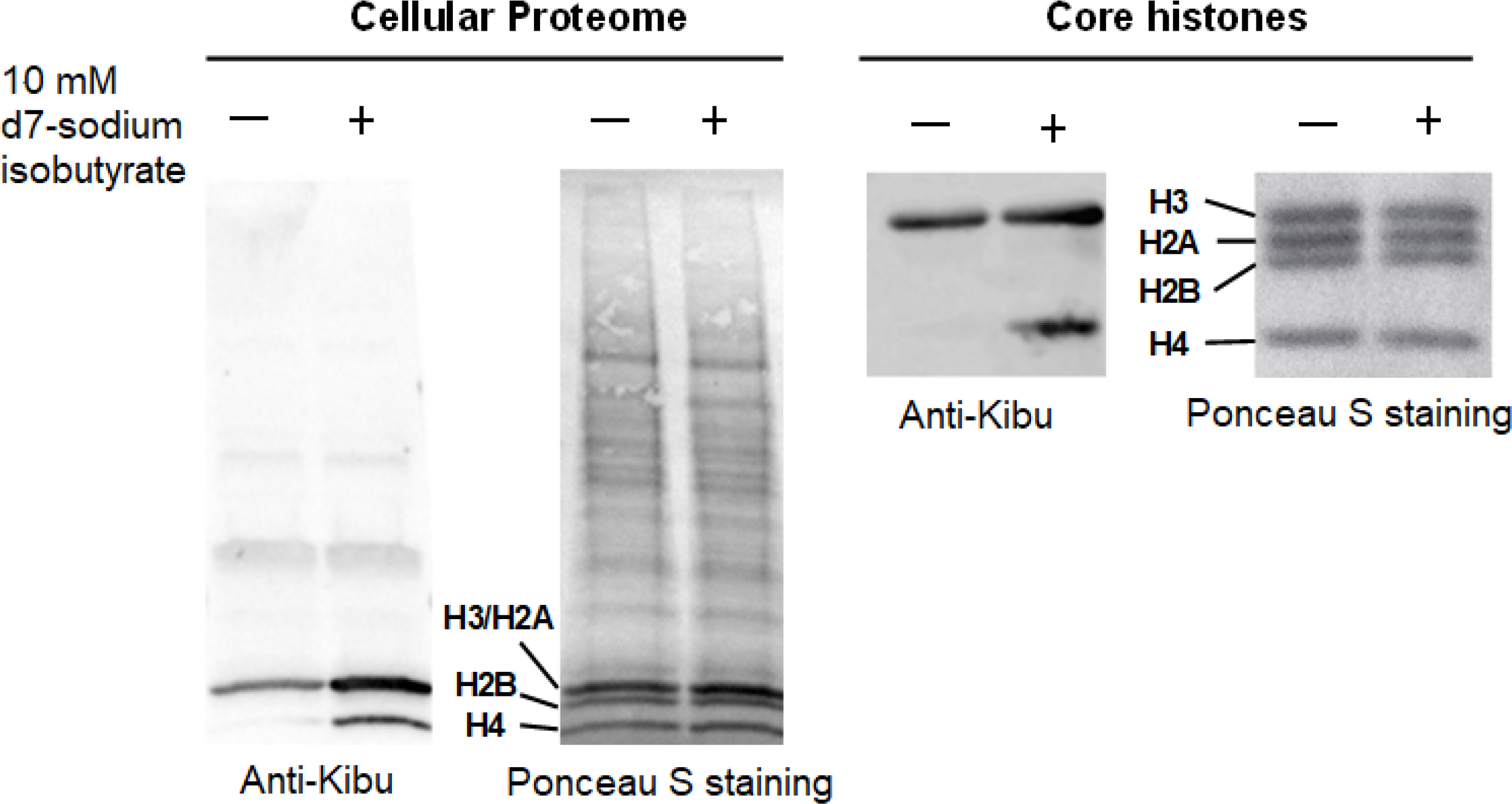
Detection of lysine isobutyrylation on protein lysines. HEK293T cells were treated with sodium isobutyrate to induce the synthesis of isobutyryl-CoA. Cellular proteome and core histone proteins were extracted and tested with anti-butyryllysine antibody (PTM Biolabs, Cat#PTM-301). Treatment of cells with isobutyrate induced increase of lysine isobutyrylation level on core histone proteins while no appreciable change was observed on non-histone protein isobutyrylation upon isobutyrate treatment.

To confirm histone Kibu in cells, we performed HPLC-MS/MS analysis of the histone extract (**Figure 6**). The core histones were extracted from the sodium d7-isobutyrate treated HEK293T cells and subsequently digested with trypsin. The resulting tryptic peptides were subjected to nano-HPLC/MS/MS analysis and protein sequence alignment with Mascot algorithm. Our analysis led to the identification of two modified H3 peptides, KSTGGKAPR and KQLATKAAR, which contain a mass shift of + 77.0858 Da at sites H3K14 and H3K23, respectively. This mass shift is the same as that caused by the addition of a d7-isobutyralation, demonstrating the existence of two novel isobutyralation sites on histone H3.

**Figure 6.**
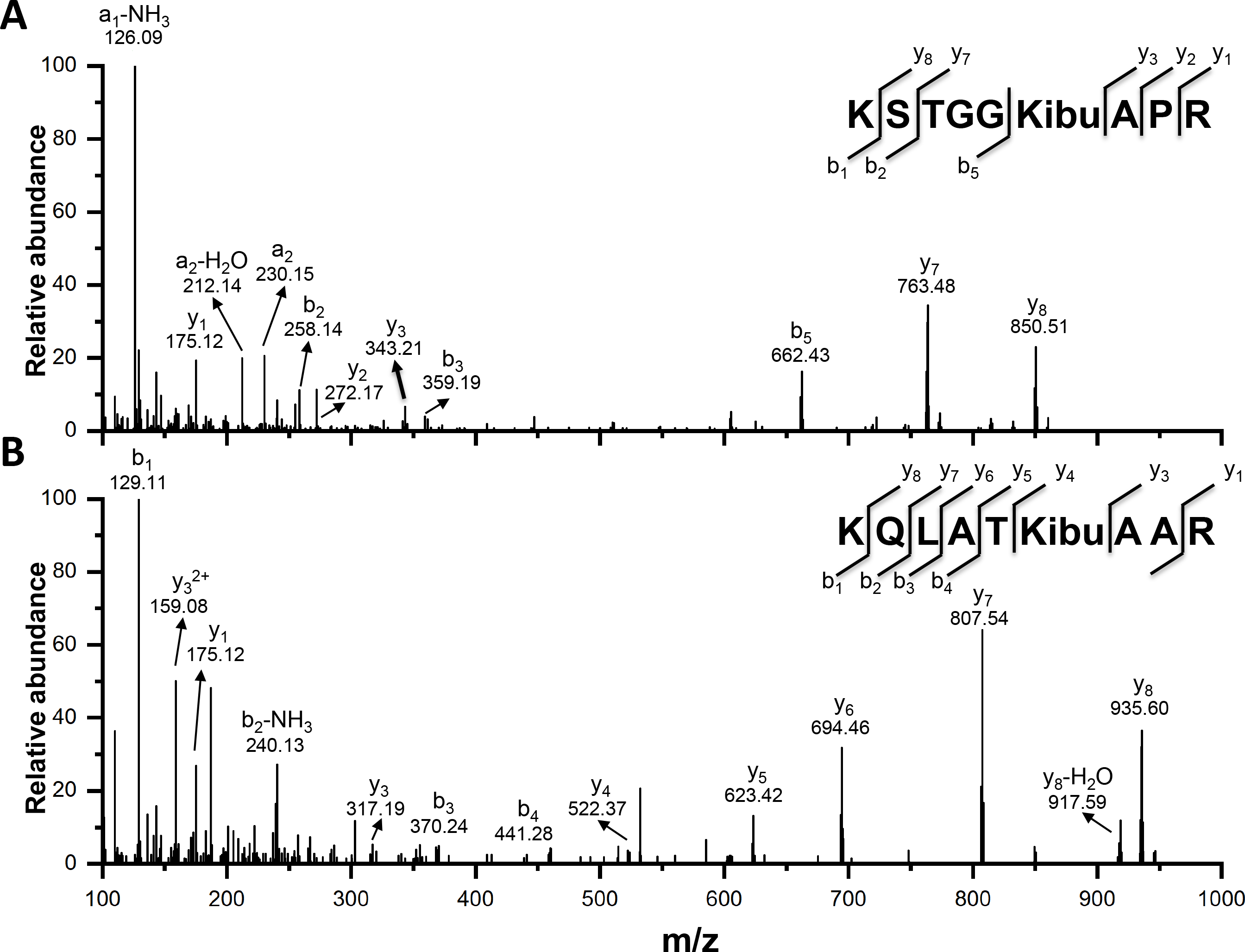
MS/MS spectra of histone H3Kibu14 (A) and H3Kibu23 (B) in 293T cells. The b and y ions refer to peptide backbone fragment ions containing N- and C-terminus, respectively.

### The p300 acetyltransferase possesses histone isobutyrylation activity in the cell

We moved on to test which KAT can mediate Kibu levels in cells. We focused on p300 and HAT1 as they both showed clear isobutyrylation activity on peptide substrates (**Figure 3** and **Figure 4**). To determine if they could catalyze *in cellulo* lysine isobutyrylation, we performed transient transfection of p300 and HAT1 plasmid in 293T cells and then treated the cells with sodium d7-isobutyrate. Consistent with the above observations (**Figure 5**), isobutyrate significantly increased Kibu levels in histones, suggesting isobutyrate is the source of isobutyryl group to promote isobutyrylation (**Supplementary Figure S4, Lanes 2 and 6**). In the presence of p300 overexpression and isobutyrate treatment together, Kibu levels were further boosted (**Supplementary Figure S4, Lane 4**) especially on histone H3 region. These data suggest that histone H3 may be a major Kibu target mediated by p300. The result is consistent with previous studies that p300 acetylates multiple sites in histone H3.^32,33^ Surprisingly, we did not observe any significant change in the Kibu level with HAT1 overexpression (**Supplementary Figure S4, Lane 8**). Such a difference between biochemical and *in cellulo* assay results in isobutyrylation may reflect the context dependence of HAT1 activity.

### Structural insights of the HAT1 and p300 interactions with isobutyryl-CoA

To elucidate the structural basis of HATs’ isobutyryltransferase activity, we generated the 1.6 Å crystal structure of the ternary complex of HAT1 with isobutyryl-CoA (IbuCoA) and histone H4(K12A) mutant peptide, referred to here as HAT1-IbuCoA-H4(K12A) (PDB ID: 6VO5) (**Table S1**). We used the H4(K12A) mutant peptide instead of the wild type counterpart to avoid any reaction intermediates between the ε-amino group of lysine12 (K12) of H4 and IbuCoA, and eventually to better resolve the electron density of the isobutyryl moiety of IbuCoA in HAT1-IbuCoA-H4(K12A). For comparison, the wild type H4K12 side chain inserts into the HAT1 active site tunnel in the crystal structure of HAT1 bound to acetyl-CoA and H4, referred to here as HAT1-AcCoA-H4 (PDB ID: 2P0W). Previous studies have shown that human HAT1 can acetylate soluble (but not nucleosomal) histone H4 at K5 and K12 positions.^34^ In HAT1-IbuCoA-H4(K12A), the H4(K12A) peptide adopts a similar conformation and binding pattern with HAT1 compared to that of HAT1-AcCoA-H4 (**Figure 7A, B**), suggesting that the N-terminal sequence motif of H4 plays a crucial role in HAT1 substrate recognition and binding, and consistent with H4K12 being the preferred HAT1 acetylation site. The overall IbuCoA interaction with HAT1 is similar to that of AcCoA-HAT1 interaction observed in HAT1-AcCoA-H4, except for the adenosine ring of IbuCoA in HAT1-IbuCoA-H4(K12A), which adopts a pi-stacking interaction with Phe288 side chain of HAT1 instead of Lys284 (**Figure 7C**). The electron density omit-map for IbuCoA in the crystal structure of HAT1-IbuCoA-H4(K12A) contained clear density for the extra isobutyryl moiety of IbuCoA, thus revealing that HAT1 also accommodates IbuCoA in its active site without any structural rearrangements (**Figure 7D**).

**Figure 7.**
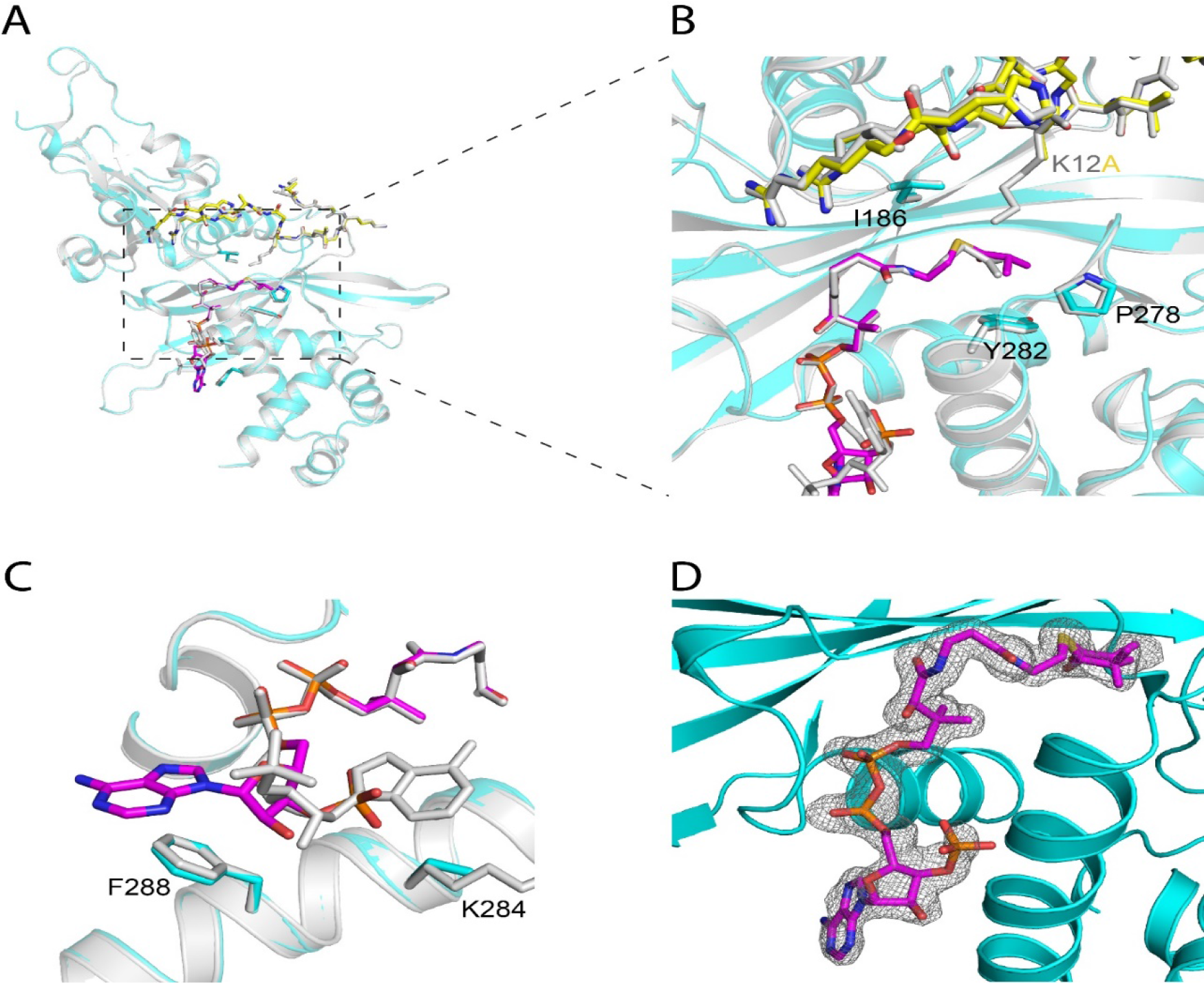
Crystal structure of HAT1 in complex with isobutyryl-CoA. (A) Overall fold of HAT1-IbuCoA-H4(K12A) shown in cyan, superposed on HAT1-AcCoA-H4 (PBD ID: 2P0W) shown in grey. IbuCoA and Histone H4(K12A) peptide is shown in stick representation in magenta in yellow, respectively. Acetyl-CoA and Histone H4 from HAT1-AcCoA-H4 structure (PBD ID: 2P0W) is shown in stick representation in grey. (B) Close-up view of the Isobutyryl moiety of IbuCoA binding site. (C) Close-up view of the adenosine ring of IbuCoA binding site. (D) The F_o_-F_c_ electron density omit-map of IbuCoA (chain B) in the crystal structure of HAT1-IbuCoA-H4(K12A), displayed as grey mesh and contoured at 2.5σ.

Similarly, p300 was previously shown to accommodate a diverse array of acyl-CoAs as substrates, and several structures of p300 in complex with acyl-CoA variants including acetyl-CoA and butyryl-CoA were reported.^19^ To further corroborate the p300 isobutyryltransferase activity observed in our assays, we modeled an IbuCoA in the crystal structure of p300 bound to butyryl-CoA (PDB ID: 5LKT). Impressively, we found that IbuCoA and butyryl-CoA bound to the active pocket of p300 with the same conformation (**Supplementary Figure S5**). Clearly p300 accommodates well isobutyryl-CoA in its acyl-CoA binding pocket, accounting for the observed Kibu activity.

### Isobutyrate globally affects the transcriptional profile of 293T cells

To systemically understand epigenetic changes affected by lysine isobutyrylation, we performed RNA-seq profiling on isobutyrate-treated 293T cells. Gene set enrichment analysis (GSEA) was used to compare the rank-ordered dataset of isobutyrate-treated versus control transcripts with respect to the KEGG pathways altered by isobutyrate treatment (**Figure 8 and supplementary Figure S6**). The results revealed that isobutyrate treatment caused upregulation of a number of genes associated with such important biological pathways as diabetes-related signaling, calcium signaling, hedgehog signaling, and JAK/STAT signaling. On the other hand, there were also some genes downregulated by isobutyrate which included aminoacyl-tRNA biosynthesis, mRNA splicing, DNA replication, and DNA repair.

**Figure 8.**
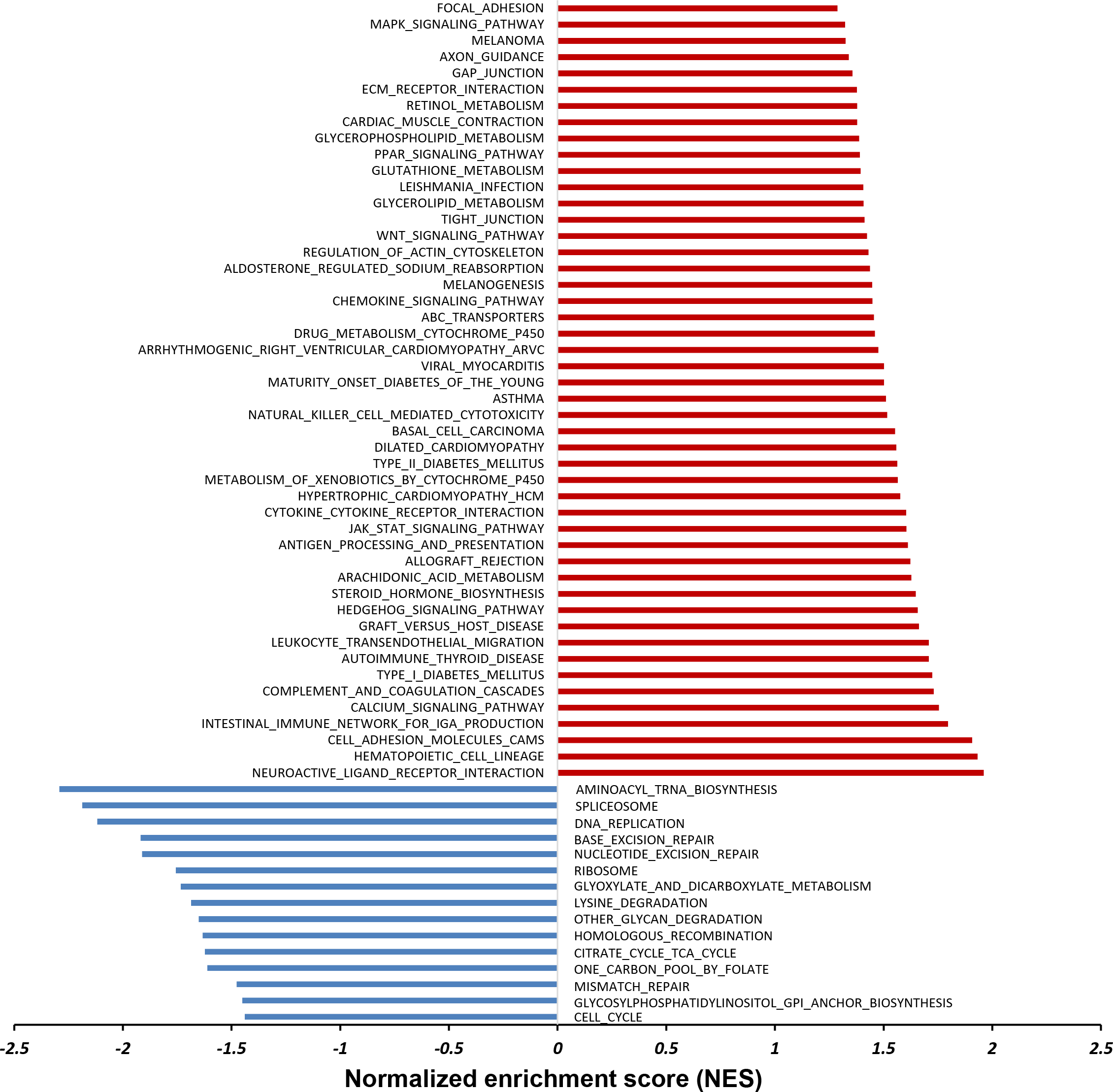
Enrichment analysis of the top gene sets altered by isobutyrate treatment in HEK293T cells. Significantly altered gene sets were defined with the False Discovery Rate (FDR) and p value less than 0.25 and 0.05, respectively. These genes were analyzed against the KEGG pathway for upregulation (red) and downregulation (blue).

## Discussion

Histone modifications are key players in the epigenetic regulation of chromatin dynamics and transcriptional programing. Various histone PTM marks in combination create unique epigenetic patterns that dictate transcriptional statuses of target genes.^35^ Lysine butyrylation was recognized as n-butyrylation upon its discovery.^18^ Later study showed that butyrylation competes with acetylation on the same lysine residue and leads to different biological outcomes.^15^ Herein, we identified lysine isobutyrylation, the structural isomer of n-butyrylation, as a new PTM in histones. Although our mass spectrometry experiment only revealed two sites in histone H3, K18 and K23, being isobutyryalted, the Western blot data suggest that histone H4 was also isobutyrylated (**Figure 5**). In future it is needed to develop isobutyryllysine-specific antibody or chemical probes and apply them to precisely map out lysine isobutyrylation sites in the chromatin histones and in the whole proteome. Both n-butyryl-CoA and isobutyryl-CoA are ample metabolites in mammalian cells but differ in their biosynthetic pathways. While n-butyryl-CoA is derived from fatty acid metabolism, isobutyryl-CoA comes from valine metabolism **(Figure 1)**. Dairy and ruminant foods are rich in branched chain fatty acids whose degradation provide another rich source of isobutyryl-CoA.^36^ Although butyryl-CoA mutase enzyme has been known in prokaryotic organisms and is able to interconvert n- and iso-butyryl-CoA,^37^ our blast search does not yield any possible orthologs existing in eukaryotic organisms. Indeed, treatment of 293T cells with d7-isobutyryate or valine only boosted isobutryl-CoA level, but not n-butyryl-CoA (**Supplementary Figure S1**), supporting that an isobutyryl-CoA mutase is not present in higher organisms.

The biochemical activity assays clearly showed that KAT members p300 and HAT1 catalyze lysine isobutyrylation activity *in vitro*. This is further substantiated by structural determination that both KATs bind to isobutyryl-CoA in their acetyl-CoA binding pocket. However only p300 showed cellular Kibu activity in our tested cell model. The discrepancy of HAT1 isobutyrylation activity in the biochemical assays from its lack of cellular activity may be a consequence of cellular context. HAT1 has been shown to form a protein complex with RIP1/3,^38^ which may change the molecular environment of HAT1 and contribute to alterations in isobutyrylation substrate specificity in cells. It will be necessary in future to investigate isobutyrylation activity of p300, HAT1, and other KATs (e.g. HBO1) in a broader scope and in different biological systems.

RNA-seq profiling showed that isobutyrate led to an extensive change in the expression levels of multiple genes in 293T cells. This exemplifies the intricate connection between metabolites and epigenetics: isobutyrate increases cellular levels of isobutyryl-CoA and thereby leads to isobutyrylation of nuclear histones and transcriptional changes. In terms of biophysical properties, lysine isobutyrylation may have similar effects as acetylation on chromatin structure and gene transcription: both acylations neutralize the positive charge on the lysine side chain and disrupt or weaken the electrostatic interaction between histones and DNA. As a result, the chromatin architecture becomes more loosely stacked and transcriptionally active gene sets are enhanced. At this point, it is unknown whether any reader protein modules exist to recognize isobutyryllysine and distinguish it from other lysine acylation marks. Our preliminary results showed that TAF1 bromodomain binds to acetylated H4 peptide but not isobutyrylated H4 peptide (data not shown), suggestive of possible antagonistic function of lysine isobutyrylation from acetylation. In addition, our RNA-seq data showed that certain genes are downregulated upon isobutyrate treatment. One reason for this phenomenon could be owing to the pharmacological effect of isobutyrate on cellular HDAC activities. Previous studies showed that n-butyrate, β-hydroxybutyrate and 4-phenylbutyrate all have inhibitory effects on the HDAC enzymes,^39^ which would positively influence gene expression by increasing the chromatin accessibility of transcriptional factors. Likely, isobutyrate may also affect the gene expression profile through inhibiting HDACs. This study calls for a comprehensive assessment of transcriptional and pharmacological effects of different short chain fatty acid (SCFA) molecules.

Acetate, propionate, and n-butyrate, produced by human gut microbiota directly impact on host physiology. ^40^ For instance, these SCFAs exhibit anti-inflammatory and anti-proliferative properties in the gastrointestinal tract, which provides novel strategies to develop anti-inflammation or anti-tumor therapeutics.^41^ It will be intriguing to examine the percentage of isobutyrate molecules produced by luminal microbiota and how it affects the physiology of the intestinal epithelial cells. Protein isobutyrylation in those host cells may dynamically changes in response to the microbiota environment. During valine catabolism, degradation of isobutyryl-CoA is catalyzed by isobutyryl-CoA dehydrogenase (IBD) encoded by the *ACAD8* gene.^42^ Mutations in *ACAD8* cause a rare inborn metabolic disorder, IBD deficiency, and lead to impaired isobutyryl-CoA breakdown and reduced energy production.^42^ Although most IBD deficiency patients are asymptomatic, some develop severe features of dilated cardiomyopathy, hypotonia, and anemia.^43^ The range of symptoms associated with IBD deficiency remain unclear, and biomarkers for IBD deficiency are being investigated to allow efficient and effective diagnosis.^44^ We anticipate that isobutyryl-CoA level would increase and hence protein lysine isobutyrylation may be upregulated in patients with IBD deficiency. Further proteomic work will be warranted to screen lysine isobutyrylation and investigate its function in IBD deficiency disease models.

## Methods

### Quantification of butyryl-CoA level in HEK293T cells

HEK-293T cells were purchased from ATCC and cultured to 90% confluence in Dulbecco’s Modified Eagle Medium (DMEM) supplemented with 10% fetal bovine serum (FBS) and 1% streptomycin-penicillin. Sodium d7-isobutyrate was prepared by adding concentrated NaOH solution to d7-isobutyric acid (Sigma-Aldrich, Cat# 632007), lyophilized and quantified by reverse phase (RP)-HPLC. Cells were treated with varied concentrations of sodium d7-isobutyrate and valine (Sigma-Aldrich, Cat#V0500) for 24 hours, and washed with ice-cold PBS buffer followed by fixing in methanol at −80 °C for 15 minutes. Cells were collected in 50% of methanol with gentle scrape, and centrifuged at 16,000 *g* at 4 °C. The supernatant was collected for HPLC-MS/MS analysis.

An Atlantis® T3 (4.6×150 mm, 3 μm) column with a Phenomenex SecurityGuard C-18 guard column (4.0 mm×2.0 mm) was applied to separate analytes. The column temperature was constant at 32 °C. The mobile phase A was 10 mM ammonium acetate, and mobile phase B was acetonitrile. A gradient method was applied for separation, with a 0.4 mL/min flow rate, (time/minute, % mobile phase B): (0, 6), (15, 30), (15.01, 100), (22.50, 100), (22.51, 6). The injection volume was 30 μL, and the autosampler injection needle was washed with methanol after each injection. Nitrogen was used as the desolvation gas at a flow rate of 500 L/h. The desolvation temperature was 500 °C and the source temperature was 120 °C. Argon was used as the collision gas, and the collision cell pressure was 3.5×10^-3^ mbar. Samples were analyzed in the positive ion mode. The capillary voltage was 3.2 kV, the cone voltage was 42 V and the collision energy was 22 eV. A multiple reaction monitoring (MRM) function was applied for the detection of analytes. The ion transition 838→331 was monitored for iso- and n-butyryl-CoA, and 845→338 for d7-isobutyryl CoA.

### Expression and purification of p300 and HAT1

The expression of p300 HAT domain (1287-1666) was done using the semisynthetic method developed by the Cole lab.^45^ A CT14 peptide aa 1653-1666 (sequence: CMLVELHTQSQDRF) was synthesized by solid-phase peptide synthesis and purified by C18 RP-HPLC. The pTYB2 plasmid encoding the inactive p300 HAT domain (region 1287-1652) fused to intein-chitin-binding domain was transformed into *E.coli* BL21(DE3)/RIL competent cells through heat-shock method and then grown on lysogeny broth (LB)-agar plates containing both ampicillin (final concentration: 100 μg/mL) and chloramphenicol (final concentration: 50 μg/mL). Colonies were harvested and grown at 37°C in 8 mL then 1 L cultures of terrific broth media containing both ampicillin and chloramphenicol. Protein expression was induced by the addition of isopropyl β-D-1-thiogalactopyranoside (IPTG) at 1.0 mM final concentration and shaken for 16 h at 16 °C. The cells were collected by centrifugation at 4000 rpm for 25 min, suspended in lysis buffer (25 mM Na-HEPES (pH 8.0), 500 mM NaCl, 1 mM MgSO_4_, 10 % glycerol, and 2 mM phenylmethanesulfonyl fluoride (PMSF)), and lysed by a microfluidic cell disruptor. The lysates were cleared by centrifugation to obtain the supernatant. Chitin resin were equilibrated with column buffer (25 mM Na-HEPES (pH 8.0), 250 mM NaCl, 1 mM EDTA, 0.1 % trition X-100, and 1 mM PMSF) before they were used to purify the supernatant. The column was thoroughly washed with column buffer and wash buffer (25 mM Na-HEPES (pH 8.0), 500 mM NaCl, 1 mM EDTA, 0.1 % triton X-100 and 1 mM PMSF) before being ligated to the CT14 peptide to obtain active p300. CT14 peptide was dissolved in cleavage buffer (25 mM Na-HEPES (pH 8.0), 250 mM NaCl, 1 mM EDTA, and 200 mM 2-mercaptoethanesulfonic acid (MESNA)) and added to the column. After the addition of the CT14 peptide in cleavage buffer, the column was shaken at room temperature for 16 h. The protein was then eluted from the column, and several volumes of cleavage buffer were added to ensure the complete elution. The eluted protein was further purified by cation exchange chromatography using the NGC fast protein liquid chromatography (FPLC) system, Biorad. The identification of p300 HAT domain was confirmed using a 12 % sodium dodecyl sulfate-polyacrylamide gel electrophoresis (SDS-PAGE). Millipore centrifugal filtration was used to concentrate protein solution and the Bradford assay was used to determine final protein concentration. Lastly, the protein was aliquoted, flash frozen by liquid nitrogen and stored at −80°C.

The expression and purification of human HAT1 (20-341) was done following the method described Hong Wu et al^46^. The pET28a-LIC-HAT1 plasmid (plasmid #25239, Addgene) was transformed into BL21 (DE3)/ RIL competent cells through heat-shock and then spread on agar plates containing antibiotics kanamycin and chloramphenicol. Protein expression was induced by the addition of IPTG (final concentration: 1.0 mM) and shaken for 16 h at 16 °C. The cells were collected and suspended in lysis buffer (50 mM Na-phosphate (pH 7.4), 250 mM NaCl, 5 mM imidazole, 5 % glycerol, 2 mM β-mercaptoethanol, and 1 mM PMSF) then disrupted using the cell disruptor. The supernatant was passed through a column containing Ni-NTA resin equilibrated with column washing buffer (20 mM Tris-HCl (pH 8.0), 250 mM NaCl, 5% glycerol, 30 mM imidazole, and 1 mM PMSF) and the resin was washed with the same column washing buffer for twice. Next, the resin was washed with buffer containing a higher concentration of imidazole (20 mM Tris-HCl (pH 8.0), 250 mM NaCl, 5% glycerol, 50 mM imidazole, and 1mM PMSF) for three times. The proteins on the resin were then eluted with elution buffer (20 mM Tris-HCl (pH 8.0), 250 mM NaCl, 5% glycerol, 500 mM imidazole, and 1 mM PMSF), and dialyzed in the dialysis buffer (25 mM Tris-HCl (pH 8.0), 150 mM NaCl, 10% glycerol, 1 mM DTT) at 4°C overnight. Thrombin was added to the dialyzed protein containing HAT1 and dialyzed in thrombin cleavage buffer (20 mM Tris-HCl (pH 8.0), 100 mM NaCl, 2.5 mM CaCl_2_, 5% glycerol, 1 mM DTT) for 20 hours at 4°C to remove the His6x-tag. The resultant protein was concentrated and purified by anion exchange chromatography using the NGC FPLC system. HAT1 purity was checked using SDS-PAGE. Millipore centrifugal filter and Bradford assay were used to concentrate and determine protein concentration, respectively. The protein was aliquoted, flash frozen by liquid nitrogen and stored at −80°C.

### Synthesis of isobutyryl-CoA

2 mmol of isobutyric acid (176.2 mg) was dissolved in 5 mL of freshly distilled CH_2_Cl_2_. To this solution was added 1 mmol of N, N’-dicyclohexylcarbodiimide (DCC) (206.4 mg), and the reaction was allowed to proceed at room temperature for 4 h. The reaction mixture was filtered to remove dicyclohexylurea (DCU), and then CH2Cl2 was removed using rotary evaporation. The dried crude material was used in the next step without further purification. 0.013 mmol of CoA hydrate (10 mg) was dissolved in 1.5 mL of 0.5 M NaHCO3 (pH 8.0) and cooled down on ice bath. Then the crude isobutyric anhydride (10.3 mg, 0.065 mmol) in 1 mL of CH3CN/acetone (1:1 v/v) was added dropwise to the CoA solution. The reaction solution was stirred at 4 °C overnight and quenched by adjusting pH to 4 with 1 M HCl. The reaction mixture was subjected to RP-HPLC purification with gradient 5–45% acetonitrile over 30 min at flow rate 5 mL/min; UV detection wavelength was fixed at 214 and 254 nm. HPLC buffer was 0.05% TFA in water (solution A) and 0.05% TFA in acetonitrile (solution B). The fractions were collected, rotary evaporated and lyophilized to yield 6.34 mg white solid. The purity of final product was checked by analytical RP-HPLC and molecular weight was confirmed by MALDI MS (found [M+H]^+^ 838.6).

### Biochemical assays of lysine acyltransferase activities of KATs

Acetyl-CoA (Sigma-Aldrich, Cat#A2181), propionyl-CoA (Chem Impex Int’l Inc, Cat# 01895) and n-butyryl-CoA (Sigma-Aldrich, Cat#B1508) salts were purchased from commercial suppliers. Synthetic histone peptides H3(1-20) or H4(1-20) (20 amino acids from the N-terminal of histone H3 and H4 (the sequence of H3(1-20) is Ac-ARTKQTARKSTGGKAPRKQL, the sequence of H4(1-20) is Ac-SGRGKGGKGLGKGGAKRHRK) were used as acyl acceptor substrates. For single-point quantification assays, 30 μM of each acyl-CoA molecule was incubated with individual KAT enzymes and 100 μM histone peptides. The enzymatic reactions were conducted in KAT reaction buffer containing 50 mM HEPES-Na and 0.1 mM EDTA-Na at pH 8.0. KAT enzymes were mixed with individual acyl-CoA molecules and peptide substrates, followed by incubation at 30 °C for 15 minutes to allow enzymatic transfer of acyl groups to lysine substrates and release of by-product CoASH. The fluorogenic probe 7-diethylamino-3-(4’-maleimidylphenyl)-4-methylcoumarin (abbr. CPM, ThermoFisher, Cat# D346) in 100% DMSO was then added to both quench the enzymatic reaction and react with CoASH to yield the fluorescent CPM-SCoA complex for fluorescence quantification ^27^. The fluorescence intensities were measured with excitation and emission wavelengths at 392 nm and 82 nm, respectively, with a FlexStation®3 microplate reader. Duplicated experiments were performed and the results were shown in a bar graph.

For kinetic characterization of p300 and HAT1 with acyl-CoA molecules, varied concentration of acyl-CoA was incubated with p300 or HAT1 and 200 μM of H4(1-20) peptide. All the reactions were conducted in the same KAT reaction buffer as the single-point assay. The fluorescence intensity was measured with the same method as the single-point assay and catalytic rate was determined based on fluorescence intensities. Kinetic constants including binding affinity (*Km*) and catalytic efficiency (*kcat*) were determined by fitting the acyl-CoA concentration-catalytic rates to the Michaelis-Menten equation using KaleidaGraph.

### Western blot analysis of HAT1 mediated *in vitro* histone H4 isobutyrylation

1 μg of human recombinant histone H4 (BioLabs, Cat# M2504S) was incubated with 50 μM n-butyryl- or isobutyryl-CoA and 0.2 μM of HAT1 at 30 °C for 1 hour. The reaction mixture was boiled in SDS-PAGE gel loading buffer and resolved on a 15% polyacrylamide gel followed by wet membrane transfer to a nitrocellulose (NC) membrane. The NC membrane was blocked with 5% non-fat milk in Tris-Buffered Saline+0.1% Tween-20 (TBST) for 1 hour at room temperature. Anti-butyryllysine antibody (PTM BioLabs, PTM#301) at 1:2000 dilution was incubated with the membrane overnight at 4 °C. The membrane was washed with TBST buffer for three times and incubated with the goat anti-rabbit IgG-HRP (Santa Cruz Biotechnology, Cat# sc-2004) with 1:3000 dilution at room temperature for 1 hour. The membrane was then washed and subjected to chemiluminescence detection with the ECL substrate (ThermoFisher, Cat# 32209) on a LI-COR Odyssey system (LI-COR Biosciences).

### Western blot analyses of *in cellulo* lysine isobutyrylation and lysine acetylation in response to isobutyrate treatment

HEK293T cells were cultured to 90% confluence in DMEM medium supplemented with 10% FBS and 1% streptomycin-penicillin antibiotics. For in-cell Kibu level analysis, cells were treated with 10 mM d7-isobutyrate for 16 hours followed by cellular protein extraction. Whole cell lysate was extracted in M-PER™ Mammalian Protein Extraction Reagent (ThermoFisher Scientific, Cat# 78501) with gentle sonication and core histone proteins were extracted with the EpiQuik Total Histone Extraction Kit (Epigentek, Cat# OP-0006-100). The extracted lysates and histone proteins were resolved on a 4-20% gradient gel (Biorad) and a 15% polyacrylamide gel, respectively. Kibu levels were detected by Western blot using the same procedure aforementionedFor in-cell Kac level analysis, cells were treated with 20 mM d7-isobutyrate for 24 hours followed by core histone extraction. Histone extracts were then resolved on a 15% polyacrylamide gel and Kac levels were determined with Western blot using anti-acetyllysine antibody (PTM Biolabs, Cat# PTM-101), with anti-Knbu/Kibu as the positive control using anti-butyryllysine antibody (PTM Biolabs, Cat# PTM-301) and histone H3 as the loading control using histone H3 antibody (Santa Cruz Biotechnology, Cat# sc-517576).

### *In cellulo* analyses of lysine isobutyrylation in response to p300 and HAT1 overexpression

HEK293T cells were cultured to 90% confluence in DMEM medium supplemented with 10% FBS and 1% streptomycin-penicillin antibiotics. Plasmids N-flag-HAT1wt (GeneCopoeia, Cat# EX-I0105-M13-11) and pCMVβ-p300-myc (Addgene, Cat# 30489) were transfected into cells with Lipofectamine 3000^TM^ Transfection Reagent (ThermoFisher, Cat# L3000008). The cells were then incubated with 20 mM sodium d7-isobutyrate for 24 hours followed by core histone extraction. Kibu levels were analyzed by Western blot using the same procedure and antibodies aforementioned.

### HAT1 crystallization and structural determination

HAT1 was expressed and purified as described previously.^46^ HAT1 at 8 mg/mL was incubated with isobutyryl coenzyme A (IBuCoA) and histone H4 K12A mutant peptide (amino acids 1–20 of H4) at a 1:10:5 molar ratio of HAT1:IbuCoA:H4(K12A) and crystallized using the sitting drop vapor diffusion method by mixing 2 μL of protein solution with 1 μL of the reservoir solution containing 2 M Sodium Dihydrogen Phosphate and 0.1 M MES, pH 6.5. Crystals were cryo-protected in the corresponding mother liquor supplemented with 30% glycerol and cryo-cooled in liquid nitrogen. X-ray diffraction date were collected at the Advanced Photon Source (APS) BEAMLINE 19-ID. Data were processed using XDS^47^ and merged with Aimless.^48^ PDB ID: 2P0W was used in Fourier transform using Refmac5.^49^ Model building and visualization was performed in COOT^50^ and the structure was validated with Molprobity.^51^ Data collection and refinement statistics are summarized in **Table S1**. The HAT1-IbuCoA-H4(K12A) structure factors and coordinates have been deposited in the Protein Data Bank with the PDB ID: 6VO5.

### HPLC/MS/MS analysis

Core histones (~4 μg) extracted from d7-sodium isobutyrate treated 293T cells were resolved in SDS-PAGE. Histones were excised from the gel, and subjected to in-gel digestion by trypsin (Tan eta al, 2011, Cell, 146: 1016-28). The digested peptide was dissolved in 2.5 μL water containing 0.1% formic acid (v/v), and then loaded onto a home-packed capillary column (10 cm length × 75 mm ID, 3 μm particle size, Dr. Maisch GmbH, Germany) which was connected to an EASY-nLC 1000 system (Thermo Fisher Scientific Inc.). The mobile phase A was water containing 0.1% formic acid (v/v), and mobile phase B was acetonitrile containing 0.1% formic acid (v/v). A 60-min gradient of 2% to 90% mobile phase B at a flow rate of 200 nl/min was used for the peptide separation. The eluted peptides were analyzed by a Q-Exactive mass spectrometer (Thermo Fisher Scientific Inc.). A Full mass scan was conducted in the Orbitrap mass analyzer in the range m/z 300 to 1,400 with a resolution of 70,000 at m/z 200. The top 15 ions were fragmented with normalized collision energy of 27 and tandem mass spectra were acquired with a mass resolution of 17,500 at m/z 200.

The obtained MS/MS spectra were searched with Mascot (Matrix Science, London, UK) against UniProt Human protein database. Mono-methylation and di-methylation on Lysine and Arginine, tri-methylation on lysine, acetylation on lysine and protein N-terminal, oxidation on methionine, and d7-isobutyrylation on lysine were specified as variable modifications. Maximum missing cleavage was set at 4, and mass tolerance was set at 10 ppm for precursor ions and ±0.05 Da for MS/MS.

### RNA-seq

Total RNA was extracted using TRIzol reagent (Thermo Fisher). Indexed libraries were constructed using the Illumina TruSeq Stranded mRNA library prep kit. Samples were then sequenced on NovaSeq (PE50) with paired-end reading. Raw reads in FASTQ files were submitted for differential expression analysis using DEseq2. Gene set enrichment analysis (GSEA) was performed against KEGG pathway. Significantly altered gene sets were defined with the False Discovery Rate (FDR) and p value less than 0.25 and 0.05, respectively.

## Acknowledgments

Y.G.Z. acknowledge the support of the National Science Foundation (grants 1507741 and 1808087), and acknowledge the Proteomics and Mass Spectrometry facility (PAMS) at UGA for the peptide MS support.

YZ is supported by supported by the University of Chicago, Nancy and Leonard Florsheim family fund (Y.Z.), NIH grants R01GM115961, R01DK118266 (Y.Z.). Y.Z. is a founder, board member, advisor to, and inventor on patents licensed to PTM Biolabs Inc (Chicago, IL) and Maponos Therapeutics Inc. (Chicago, IL).

ML is supported by NIH grants R35GM131858 and 5P30CA0087. ML is a member of the Scientific Advisory Board for Epi One Inc.

We thank Aiping Dong for reviewing and depositing the HAT1 structure. Structural results shown in this report are derived from work performed at Argonne National Laboratory, Structural Biology Center (SBC) at the Advanced Photon Source. SBC-CAT is operated by UChicago Argonne, LLC, for the U.S. Department of Energy, Office of Biological and Environmental Research under contract DE-AC02-06CH11357. The Structural Genomics Consortium is a registered charity (no: 1097737) that receives funds from AbbVie; Bayer Pharma AG; Boehringer Ingelheim; Canada Foundation for Innovation; Eshelman Institute for Innovation; Genome Canada through Ontario Genomics Institute [OGI-055]; Innovative Medicines Initiative (EU/EFPIA) [ULTRA-DD: 115766]; Janssen, Merck & Co.; Novartis Pharma AG; Ontario Ministry of Research Innovation and Science (MRIS); Pfizer, São Paulo Research Foundation-FAPESP, Takeda and the Wellcome Trust.

## References

1. Aebersold, R. et al. How many human proteoforms are there? Nat Chem Biol 14, 206–214 (2018).

2. Sabari, B.R. et al. Intracellular crotonyl-CoA stimulates transcription through p300-catalyzed histone crotonylation. Mol Cell 58, 203–15 (2015).

3. Verdin, E. & Ott, M. 50 years of protein acetylation: from gene regulation to epigenetics, metabolism and beyond. Nat Rev Mol Cell Biol 16, 258–64 (2015).

4. Barnes, C.E., English, D.M. & Cowley, S.M. Acetylation & Co: an expanding repertoire of histone acylations regulates chromatin and transcription. Essays Biochem 63, 97–107 (2019).

5. Hirschey, M.D. & Zhao, Y. Metabolic Regulation by Lysine Malonylation, Succinylation, and Glutarylation. Mol Cell Proteomics 14, 2308–15 (2015).

6. Sabari, B.R., Zhang, D., Allis, C.D. & Zhao, Y. Metabolic regulation of gene expression through histone acylations. Nat Rev Mol Cell Biol 18, 90–101 (2017).

7. Lin, H., Su, X. & He, B. Protein lysine acylation and cysteine succination by intermediates of energy metabolism. ACS Chem Biol 7, 947–60 (2012).

8. Bos, J. & Muir, T.W. A Chemical Probe for Protein Crotonylation. J Am Chem Soc 140, 4757–4760 (2018).

9. He, M., Han, Z., Liu, L. & Zheng, Y.G. Chemical Biology Approaches for Investigating the Functions of Lysine Acetyltransferases. Angew Chem Int Ed Engl 57, 1162–1184 (2018).

10. Marmorstein, R. & Zhou, M.M. Writers and readers of histone acetylation: structure, mechanism, and inhibition. Cold Spring Harb Perspect Biol 6, a018762 (2014).

11. Bheda, P., Jing, H., Wolberger, C. & Lin, H. The Substrate Specificity of Sirtuins. Annu Rev Biochem 85, 405–29 (2016).

12. Schneider, A. et al. Acetyltransferases (HATs) as Targets for Neurological Therapeutics. Neurotherapeutics 10, 568–588 (2013).

13. Di Cerbo, V. & Schneider, R. Cancers with wrong HATs: the impact of acetylation. Brief Funct Genomics 12, 231–43 (2013).

14. Mohammad, H.P., Barbash, O. & Creasy, C.L. Targeting epigenetic modifications in cancer therapy: erasing the roadmap to cancer. Nat Med 25, 403–418 (2019).

15. Goudarzi, A. et al. Dynamic Competing Histone H4 K5K8 Acetylation and Butyrylation Are Hallmarks of Highly Active Gene Promoters. Mol Cell 62, 169–80 (2016).

16. Wu, Z. et al. A chemical proteomics approach for global analysis of lysine monomethylome profiling. Mol Cell Proteomics 14, 329–39 (2015).

17. Wang, X., Sidoli, S., Garcia, B. A. Application of Mass Spectrometry in Translational Epigenetics. Epigenetic Technological Applications, 55–78 (2015).

18. Chen, Y. et al. Lysine propionylation and butyrylation are novel post-translational modifications in histones. Mol Cell Proteomics 6, 812–9 (2007).

19. Kaczmarska, Z. et al. Structure of p300 in complex with acyl-CoA variants. Nat Chem Biol 13, 21–29 (2017).

20. Robinson, W.G., Nagle, R., Bachhawat, B.K., Kupiecki, F.P. & Coon, M.J. Coenzyme-a Thiol Esters of Isobutyric, Methacrylic, and Beta-Hydroxyisobutyric Acids as Intermediates in the Enzymatic Degradation of Valine. J Biol Chem 224, 1–11 (1957).

21. Yun, J.W. et al. A novel ACAD8 mutation in asymptomatic patients with isobutyryl-CoA dehydrogenase deficiency and a review of the ACAD8 mutation spectrum. Clin Genet 87, 196–8 (2015).

22. Santra, S., Macdonald, A., Preece, M.A., Olsen, R.K. & Andresen, B.S. Long-term outcome of isobutyryl-CoA dehydrogenase deficiency diagnosed following an episode of ketotic hypoglycaemia. Mol Genet Metab Rep 10, 28–30 (2017).

23. Grevengoed, T.J., Klett, E.L. & Coleman, R.A. Acyl-CoA metabolism and partitioning. Annu Rev Nutr 34, 1–30 (2014).

24. Abdinejad, A., Fisher, A.M. & Kumar, S. Production and utilization of butyryl-CoA by fatty acid synthetase from mammalian tissues. Arch Biochem Biophys 208, 135–45 (1981).

25. Webster, L.T., Jr., Gerowin, L.D. & Rakita, L. Purification and Characteristics of a Butyryl Coenzyme a Synthetase from Bovine Heart Mitochondria. J Biol Chem 240, 29–33 (1965).

26. Roe, C.R. et al. Isolated isobutyryl-CoA dehydrogenase deficiency: an unrecognized defect in human valine metabolism. Mol Genet Metab 65, 264–71 (1998).

27. Gao, T., Yang, C. & Zheng, Y.G. Comparative studies of thiol-sensitive fluorogenic probes for HAT assays. Anal Bioanal Chem 405, 1361–71 (2013).

28. Han, Z. et al. Revealing the protein propionylation activity of the histone acetyltransferase MOF (males absent on the first). J Biol Chem 293, 3410–3420 (2018).

29. Yang, C. et al. Labeling lysine acetyltransferase substrates with engineered enzymes and functionalized cofactor surrogates. J Am Chem Soc 135, 7791–4 (2013).

30. Yang, Y.Y., Ascano, J.M. & Hang, H.C. Bioorthogonal chemical reporters for monitoring protein acetylation. J Am Chem Soc 132, 3640–1 (2010).

31. Pougovkina, O., te Brinke, H., Wanders, R.J.A., Houten, S.M. & de Boer, V.C.J. Aberrant protein acylation is a common observation in inborn errors of acyl-CoA metabolism. J Inherit Metab Dis 37, 709–714 (2014).

32. Henry, R.A., Kuo, Y.M., Bhattacharjee, V., Yen, T.J. & Andrews, A.J. Changing the selectivity of p300 by acetyl-CoA modulation of histone acetylation. ACS Chem Biol 10, 146–56 (2015).

33. Dancy, B.M. & Cole, P.A. Protein lysine acetylation by p300/CBP. Chem Rev 115, 2419–52 (2015).

34. Verreault, A., Kaufman, P.D., Kobayashi, R. & Stillman, B. Nucleosomal DNA regulates the core-histone-binding subunit of the human Hat1 acetyltransferase. Curr Biol 8, 96–108 (1998).

35. Papait, R. et al. Genome-wide analysis of histone marks identifying an epigenetic signature of promoters and enhancers underlying cardiac hypertrophy. Proc Natl Acad Sci U S A 110, 20164 (2013).

36. Ran-Ressler, R.R., Bae, S., Lawrence, P., Wang, D.H. & Brenna, J.T. Branched-chain fatty acid content of foods and estimated intake in the USA. Br J Nutr 112, 565–72 (2014).

37. Jost, M., Born, D.A., Cracan, V., Banerjee, R. & Drennan, C.L. Structural Basis for Substrate Specificity in Adenosylcobalamin-dependent Isobutyryl-CoA Mutase and Related Acyl-CoA Mutases. J Biol Chem 290, 26882–98 (2015).

38. Carafa, V. et al. RIP1-HAT1-SIRT Complex Identification and Targeting in Treatment and Prevention of Cancer. Clin Cancer Res 24, 2886–2900 (2018).

39. Chriett, S. et al. Prominent action of butyrate over β-hydroxybutyrate as histone deacetylase inhibitor, transcriptional modulator and anti-inflammatory molecule. Sci Rep 9, 742 (2019).

40. Louis, P., Hold, G.L. & Flint, H.J. The gut microbiota, bacterial metabolites and colorectal cancer. Nat Rev Microbiol 12, 661–72 (2014).

41. Parada Venegas, D. et al. Short Chain Fatty Acids (SCFAs)-Mediated Gut Epithelial and Immune Regulation and Its Relevance for Inflammatory Bowel Diseases. Front Immunol 10, 277 (2019).

42. Nguyen, T.V. et al. Identification of isobutyryl-CoA dehydrogenase and its deficiency in humans. Mol Genet Metab 77, 68–79 (2002).

43. Santra, S., Macdonald, A., Preece, M.A., Olsen, R.K. & Andresen, B.S. Long-term outcome of isobutyryl-CoA dehydrogenase deficiency diagnosed following an episode of ketotic hypoglycaemia. Mol Genet Metab Rep 10, 28–30 (2017).

44. Oglesbee, D. et al. Development of a newborn screening follow-up algorithm for the diagnosis of isobutyryl-CoA dehydrogenase deficiency. Genet Med 9, 108–116 (2007).

45. Thompson, P.R. et al. Regulation of the p300 HAT domain via a novel activation loop. Nat Struct Mol Biol 11, 308–15 (2004).

46. Wu, H. et al. Structural basis for substrate specificity and catalysis of human histone acetyltransferase 1. Proc Natl Acad Sci U S A 109, 8925–30 (2012).

47. Kabsch, W. Xds. Acta Crystallogr D Biol Crystallogr 66, 125–32 (2010).

48. Winn, M.D. et al. Overview of the CCP4 suite and current developments. Acta Crystallogr D Biol Crystallogr 67, 235–42 (2011).

49. Evans, P.R. & Murshudov, G.N. How good are my data and what is the resolution? Acta Crystallogr D Biol Crystallogr 69, 1204–14 (2013).

50. Emsley, P., Lohkamp, B., Scott, W.G. & Cowtan, K. Features and development of Coot. Acta Crystallogr D Biol Crystallogr 66, 486–501 (2010).

51. Williams, C.J. et al. MolProbity: More and better reference data for improved all-atom structure validation. Protein Sci 27, 293–315 (2018).

